# Composition of Carotid Plaques Differs Between Chinese and United States Patients: A Histology Study

**DOI:** 10.1101/2025.06.10.658993

**Authors:** Jingli Cao, Marina Ferguson, Jie Sun, Mi Shen, Randy Small, Daniel S. Hippe, Xihai Zhao, Dong Zhang, Hiroko Watase, Chun Yuan, Peiyi Gao, James Kevin DeMarco, Roberto F. Nicosia, Yajie Wang, Haowen Li, Zirui Li, Yi Wang, Ted Kohler, Thomas Hatsukami, Binbin Sui

## Abstract

**Background:** The clinical manifestations of cerebrovascular disease are known to differ between the Chinese and United States (U.S.) populations as do the plaque features on imaging.

**Objectives:** The aim of this study was to investigate and compare the histological features of excised carotid plaques from Chinese and U.S. patients.

**Methods:** Carotid endarterectomy specimens collected from two prospective studies were included. The entire plaque was serially sectioned (10 µm thickness) at 0.5-1 mm intervals. Hematoxylin and eosin staining and Mallory’s trichrome staining were performed. The morphology and components of the plaques were measured and compared between the two groups.

**Results:** A total of 1,152 histological sections from 75 Chinese patients and 1,843 sections from 111 U.S. patients were analyzed. The Chinese group had significantly smaller minimum lumen diameters (median: 1.1 vs. 1.3 mm, p=0.046) and a larger percent wall volume (median: 74% vs. 70%, p=0.018) than the U.S. group. After adjusting for confounding factors, carotid plaques in the Chinese population were more likely to have more lipid pools (β=10.0%, 95%CI: 4.9 to 15.9%), more recent intraplaque hemorrhage (IPH; β=8.4%, 95%CI: 4.5 to 12.7%), and less late IPH (β=-8.2%, 95%CI: −11.3 to −5.4), and fewer fibrous cap disruptions (45% vs. 67%, p=0.061). Chinese plaques were more homogeneous and had a higher percentage of plaques with features of xanthomas than did U.S. plaques (20% vs 2.7%, p<0.001).

**Conclusions:** The histology of Chinese plaques differs significantly from that of U.S. plaques, suggesting substantial differences in the pathophysiology of atherosclerotic cerebrovascular disease between Chinese and North American populations, which could enhance the gap in racial pathology comparison, indicating a need for a different management approach.

**Central illustration:** 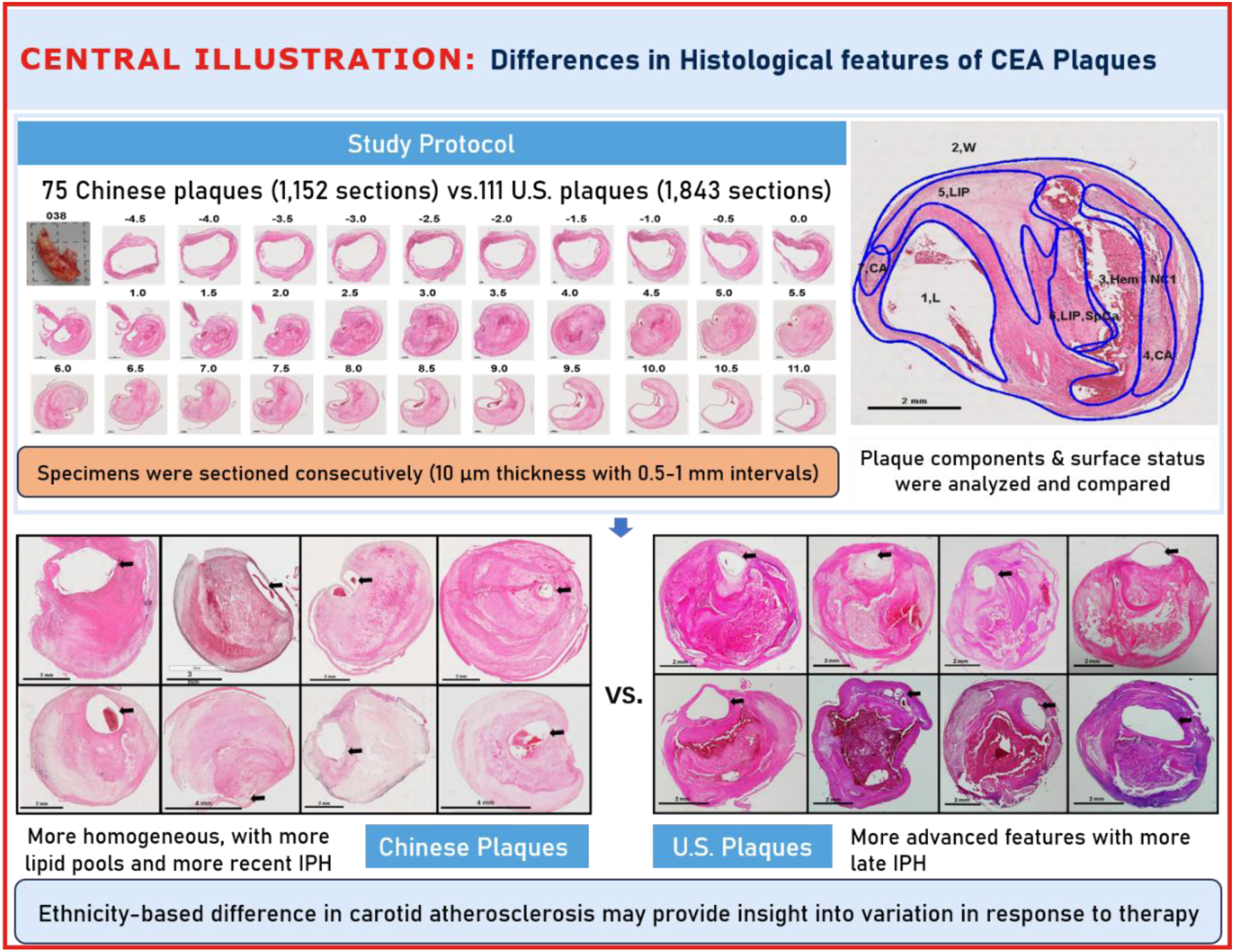

## Introduction

Stroke is one of the leading causes of death and serious long-term disability in China and the United States.^1–3^ While these populations share the burden of this potentially preventable disease, there appear to be important differences in the pattern of cerebrovascular disease between China and the U.S. When considering atherosclerosis of the large arteries, extracranial disease is more prominent in Caucasian than in Asian patients. Conversely, there is up to a five-fold higher prevalence of intracranial plaque among Chinese patients compared to Caucasians.^4, 5^ There is also a difference in the distribution of culprit lesions, which are responsible for ischemic events associated with plaque rupture. Intracranial plaques with culprit lesion features are seen three times more frequently in China than in the U.S.^6^ Our previous pilot studies using magnetic resonance imaging (MRI) also found striking differences in the location and characteristics of potential culprit plaques between two racial groups.^7,8^

Ethnicity-based characteristics and differences have been highlighted in previous studies on coronary, intracranial artery, and carotid intima-media thickness.^9–11^ Difference in the characteristics of atherosclerotic disease may be linked to different approaches to treatment and prevention. However, the histological differences in carotid advanced atherosclerotic plaques have not been extensively investigated.

Herein we report a study comparing the pathology of excised carotid plaques from patients undergoing carotid endarterectomy (CEA) in the U.S. and China. We quantified and compared plaque components of endarterectomy specimens from the University of Washington (UW) and from the Beijing Tiantan Hospital (BTH). Plaques were removed intact to preserve luminal surface morphology and were comprehensively analyzed in toto.

## Methods

### Study Population

CEA specimens were collected from 75 BTH patients in the Culprit Plaque in Acute Cerebral infarction (CPAC) study between 2014 and 2018 and from 111 patients in the UW study between 1997 and 2008. Less than 3% of the subjects enrolled in the U.S. cohort have been Asian, which reflects the prevalence of Asian-American patients undergoing CEA in the UW study. The indications for operation in both countries followed the guidelines of the Asymptomatic Carotid Atherosclerosis Study,^12^ the European Carotid Surgery Trial,^13^ and the North American Symptomatic Carotid Endarterectomy Trial.^14^ Clinical risk factors and medication use (statins, antiplatelet agents, and anticoagulant agents) were recorded through retrospective review of the medical chart. Approval was obtained from the local Institutional Research Boards at each institution (UW STUDY 00003635, BTW KY2014-004-01), and the procedures followed were in accordance with institutional guidelines. All subjects gave written informed consent.

### Surgery and Specimen Processing

All specimens were removed without disruption by incising the carotid longitudinally through the adventitia and outer media but not into the plaque. Thus, the plaque wall and luminal surface were never violated (Supplement figure 1, Right). The thickness of the artery wall left behind was less than 1 mm.

Protocols for specimen processing were standardized between the two centers. Plaques were processed by two histologists: Marina Ferguson, MSF, with over 20-years’ experience in plaque histological analysis and Jingli Cao, JLC, with over 10-years’ experience. First, the plaques were gently flushed with Ringer’s lactate and placed in 10% neutral buffered formalin for a minimum of 24 hours. Specimens were decalcified with RapidCal·ImunoTM (Zhongshan GoldenBridge Biotechnology, Beijing, China) in China and 10% formic acid in the US. Specimens were embedded en bloc in paraffin and sectioned (10 µm thickness) at 1 mm intervals in the common and 0.5 mm intervals in the internal carotid artery. The entire specimen was processed and analyzed. The sections were stained with hematoxylin and eosin (H&E) and with Mallory’s trichrome.

### Plaque Morphometry Analysis

Images for the Chinese specimens were captured on Aperio Image Scope, Pathology Viewing System v12.0.1.5027 (Leica Biosystems, Wetzler, Germany) and Ventana Image Viewer v.3.1.3 (Ventana Medical Systems, Tucson, Arizona). U.S. images were captured with a Nikon Coolpix Camera. Images of H&E sections at 1 x magnification were transferred to MATLAB R2014a (MathWorks, Natick, Massachusetts) for digitizing with QVAS (Quantitative Vascular Analysis System, 2004 Histology Version, Vascular Imaging Laboratory, University of Washington, Seattle).^15^ During digitizing, reference was made to the original glass slides using 200 x magnification to classify plaque features based on detailed analysis of their components.

All histopathological images were analyzed by a single reviewer (Marina Ferguson, MSF). For each cross-sectional location, the total area of the specimen was measured by outlining its outer border, and the lumen area by the inner perimeter of the wall. The wall area was calculated as the difference between the total area and the lumen area. Total vessel volume and wall volume were calculated by summing the corresponding areas and then multiplying by the spacing interval (0.5 or 1.0 mm). Overall plaque burden was defined as 100% x wall volume/total vessel volume. The percent narrowing at each section was calculated as 100% x (1 – lumen area/total vessel area), and the maximum value was recorded as the stenosis degree.

### Classification and Analysis of Plaque Components and Surface Status

Histologic features of endarterectomy specimens from the two groups were identified, classified and outlined. For each component, the corresponding percentages of component volume was calculated as: 100% x component volume/wall volume. Plaque compositional features were classified according to the histopathologic system described by Stary et al, from the Committee on Vascular Lesions of the Council of Atherosclerosis, American Heart Association.^15^ These were as follows (Supplement Figure 2): 1) **Lipid pools** are early lesions of morphologically distinct spaces composed of phospholipids, cholesterol monohydrate and cholesterol esters. Due to the extraction of fats from the plaque by alcohol during histological processing, lipid pools are visualized by H&E as acellular spaces that may contain small cholesterol clefts and faintly staining, grainy, amorphous material.^15^ Lipid pools do not stain with Mallory’s Trichrome, which stains fibrin, or anti-glycophorin A, which attaches to the sialoglycoprotein glycophorin A on the red cell membrane.^16^ 2) **Necrotic core** is a region filled with necrotic debris and which displace fibrous tissue. The core may have a well-defined edge populated by an inflammatory response. It may also contain lipid, cholesterol clefts, red blood cell fragments, collagen fragments, and calcifications.^17^ Regions containing a necrotic core were distinguished from areas of dense, collagen-rich fibrous intimal tissue and regions with loose connective tissue matrix. 3) **Recent intraplaque hemorrhage** is a mixture of intact and degenerating red blood cells, hemosiderin-laden macrophages, giant cells, and cholesterol crystals. Peripheral neoangiogenesis and speckled calcifications also may be present.^18^ Red blood cells are eosinophilic and lose some of this tintorial quality when they degenerate and become dehemoglobinized. Therefore, hemorrhagic areas show various degrees of pink staining on hematoxylin and eosin. Mallory’s trichrome results in a red color for fibrin, fibrin products and red blood cells, whereas extracellular matrix components stain blue. When hemorrhage occurs outside a necrotic core it is composed of intact and decomposing red blood cells, fibrin and fibrin products and a few inflammatory cells. 4) **Late intraplaque hemorrhage** is an amorphous, faintly eosinophilic material which may contain small and large cholesterol clefts along with cellular debris, hemosiderin, and a variety of calcifications.^18^ The hemorrhagic nature of this material, which no longer contains intact red blood cells, is confirmed by staining for the red blood cell membrane protein glycophorin A which provides specific evidence of prior bleeding in these areas.^19^ 5) **Calcification** is composed of hydroxyapatite crystals ranging in size from small intracellular granules^20^ to large amorphous extracellular deposits.^21^ The decalcification process removes enough calcium to permit histological sectioning, but traces of calcium, bound to phosphate or carbonate anions, remain on the periphery of the calcified region giving a characteristic blue outline when stained with hematoxylin. 6) **Loose Extracellular Matrix** is characterized by interlacing, loosely organized collagen fibrils in an amorphous proteoglycan-rich background which is populated by smooth muscle cells, fibroblasts and varying numbers of microvessels. A similar type of matrix is found in organizing thrombi, or on the luminal surface. 7) **Xanthomas** are distinct accumulations of lipid laden macrophages (foam cells) separated by richly vascularized collagenous matrix and arranged in a lobular pattern. A focal mixed inflammatory infiltrate is frequently present. Cholesterol clefts are sometimes noted. Xanthomas do not have the lipid rich necrotic center that is typically present in the fibroatheromas where foam cells are haphazardly arranged and randomly admixed with cholesterol clefts and cell debris. Xanthomas demonstrate delicate interstitial collagen and are similar to those found in patients with lipid disorders (Fig. 1).^22, 23^

**Figure 1.**
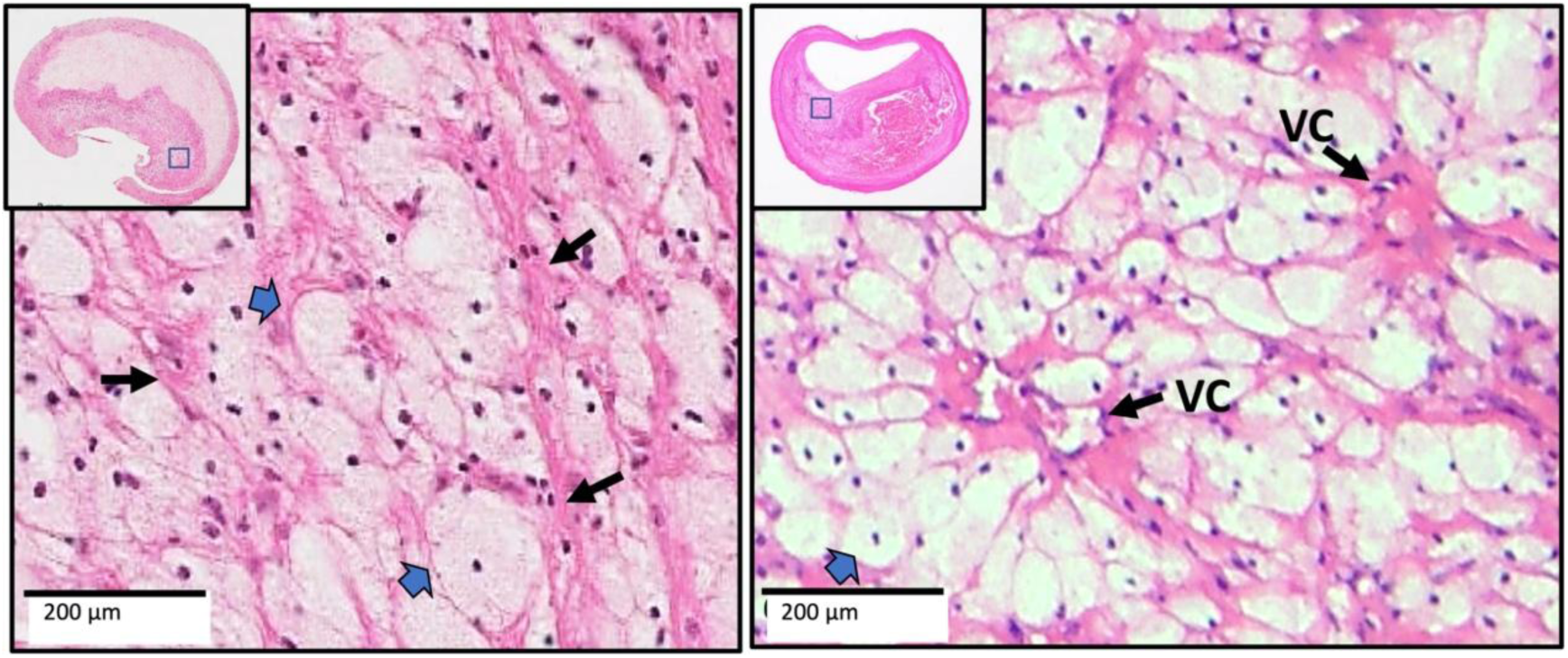
Two xanthomas with similar characteristics are shown from China (left) and from the US (right). Narrow arrows point to the delicate interstitial collagen fibrils, and block arrows point to foamy macrophages. The arrangement of macrophages around vascular channels (VC) is characteristic of xanthomas. (H&E)

Plaque surface disruptions were defined and characterized as follows: 1) **Fibrous cap disruption**: a disruption of the fibrous tissue between the necrotic core and the lumen; 2) **Disruption with mural thrombus**: a lumen surface lacking an endothelial cell covering and with an attached thrombus composed of blood components such as platelets, fibrin, red and white blood cells; 3) **Disruption with juxtaluminal calcification**: calcifications protruding into the lumen from a luminal plaque disruption.

### Data Analysis

All data were tested for normal distribution. Continuous variables and summarized as median (inter-quartile range [IQR]) and categorical variables were summarized as count (percentage). The Wilcoxon rank-sum test and Fisher’s exact test were used to compare clinical and plaque characteristics between two groups. Plaque composition and luminal surface status variables were also compared between two groups using linear and logistic regression, respectively, each with adjustments for recent carotid symptoms, carotid stenosis, sex, age, and other cardiovascular risk factors and medication. The linear regression coefficient (β) and the adjusted odds ratio (OR) with 95%CI were calculated.

Associations of plaque composition and surface conditions with recent carotid symptoms were assessed using logistic regression models with symptoms as the outcome variable. The square root transformation was applied to each composition volume variable to reduce skewness before inclusion in the corresponding model. Each model was adjusted for cohort (Chinese vs. U.S.), carotid stenosis, sex, age, and the other factors listed in Table 1. Differences in the associations with symptoms between Chinese and U.S. groups (summarized using the OR) were tested by adding a cohort-by-plaque-feature interaction term to each model. All statistical calculations were conducted with the statistical computing language R (version 4.0.3; R Foundation for Statistical Computing, Vienna, Austria).

**Table 1.** Demographic data and Clinical characteristics of the patients.

|  | Cohort* |  | P-value† |
| --- | --- | --- | --- |
|  | Chinese group<br>(N=75) | U.S. group<br>(N=111) |  |
| CVD Risk Factors |  |  |  |
| Male sex | 58 (77.3) | 101 (91.0) | 0.011 |
| Age, years | 63 (58 - 67) | 70 (62 - 76) | <0.001 |
| Hypertension | 50 (66.7) | 91 (82.0) | 0.023 |
| Hyperlipidemia‡ | 29 (39.7) | 90 (85.7) | <0.001 |
| Diabetes | 24 (32.0) | 28 (25.2) | 0.32 |
| Current smoker | 27 (36.0) | 41 (36.9) | >0.99 |
| CVD History |  |  |  |
| Symptomatic carotid artery | 22 (29.3) | 37 (33.3) | 0.63 |
| Coronary artery disease | 10 (13.3) | 50 (45.0) | <0.001 |
| Medications |  |  |  |
| Statins | 24 (32.0) | 74 (66.7) | <0.001 |
| Antiplatelets or anticoagulants | 15 (20.0) | 75 (67.6) | <0.001 |

\*Values no. (%) or median (range).
†Fisher's exact test or the Wilcoxon rank-sum test.
‡Eight subjects were missing a history of hyperlipidemia (6 Chinese and 2 U.S.).

## Results

### Study Population and Vascular Risk Factors

A total of 1,152 sections from 75 Chinese plaques and 1,843 sections from 111 U.S. plaques were analyzed. The number of sections per plaque were similar (Chinese group: median 15, range 5-28; U.S. group: median 17, range 6-34). The demographic data of the Chinese and U.S. groups are listed in Table 1. The Chinese group was younger with more females and less hypertension, hyperlipidemia, and coronary artery disease (all p<0.05). No significant difference was found in the proportion of symptomatic carotids between the two cohorts (29% vs. 33%, p=0.63). Significantly fewer Chinese patients were taking statins or anti-thrombotics prior to admission (all p<0.001).

### Plaque Morphology

The maximum outer wall diameters were similar between the two groups (median: 9.2 vs. 9.2 mm, p=0.82), as were the plaque lengths (median: 14 vs. 15 mm, p=0.66) (Table 2). The Chinese group had significantly smaller minimum lumen diameters than the U.S. group (median: 1.1 vs. 1.3 mm, p=0.046), which translated into a small though statistically significantly more severe stenosis in the Chinese plaques (median [IQR]: 97% [93 - 99%] vs. 96% [91 - 98%], p=0.005). The Chinese plaques also had a larger overall plaque burden than the U.S. plaques in terms of percent wall volume (median: 74% vs. 70%, p=0.018).

**Table 2.** Comparison of plaque morphology between two groups.

| Morphology | Cohort* |  | P-value† |
| --- | --- | --- | --- |
|  | Chinese groups<br>(N=75) | U.S. group<br>(N=111) |  |
| Plaque length, mm | 14 (10 - 17) | 15 (11 - 18) | 0.66 |
| Maximum outer wall diameter, mm | 9.2 (8.2 - 10.0) | 9.2 (8.3 - 10.0) | 0.82 |
| Minimum lumen diameter, mm | 1.1 (0.6 - 1.6) | 1.3 (0.9 - 1.9) | 0.046 |
| Stenosis, % | 97 (93 - 99) | 96 (91 - 98) | 0.005 |
| Percent wall volume, % | 74 (66 - 81) | 70 (64 - 76) | 0.018 |

\*Values no. (%) or median (inter-quartile range).
†Fisher's exact test or the Wilcoxon rank-sum test.

### Plaque Composition

When examined at low magnification, Chinese plaques had a more homogeneous morphology (Fig. 2) than U.S. plaques, which were frequently complex due to a greater admixture of different components (Fig. 3). Plaques from the U.S. cohort were composed of lesions typical of type VI atheromas where most IPH spills into a necrotic core. Features that contributed to the complexity of the U.S. plaques included broader areas of necrosis, calcifications, thrombosis, and hemorrhage. Chinese specimens contained large lipid deposits, necrotic debris, cholesterol clefts, and uniform extracellular matrix. The plaques had prominent fibro-lipid accumulations with foam cells sequestered along fibers of loose extracellular matrix (Fig. 4). In Chinese plaques, old IPH was found in necrotic cores, but additional recent IPH were often found in fibrous matrix or adjacent to xanthomas or inflammatory infiltrate (Fig. 5). Chinese plaques more frequently had lipid in foam cells with the pattern of xanthomas (15/75, 20%) than in Western plaques (3/111, 2.7% p<0.001). All components except late hemorrhage were significantly correlated with percent stenosis, and all features except calcification were significantly associated with percent wall volume (Supplement Table 1).

**Figure 2.**
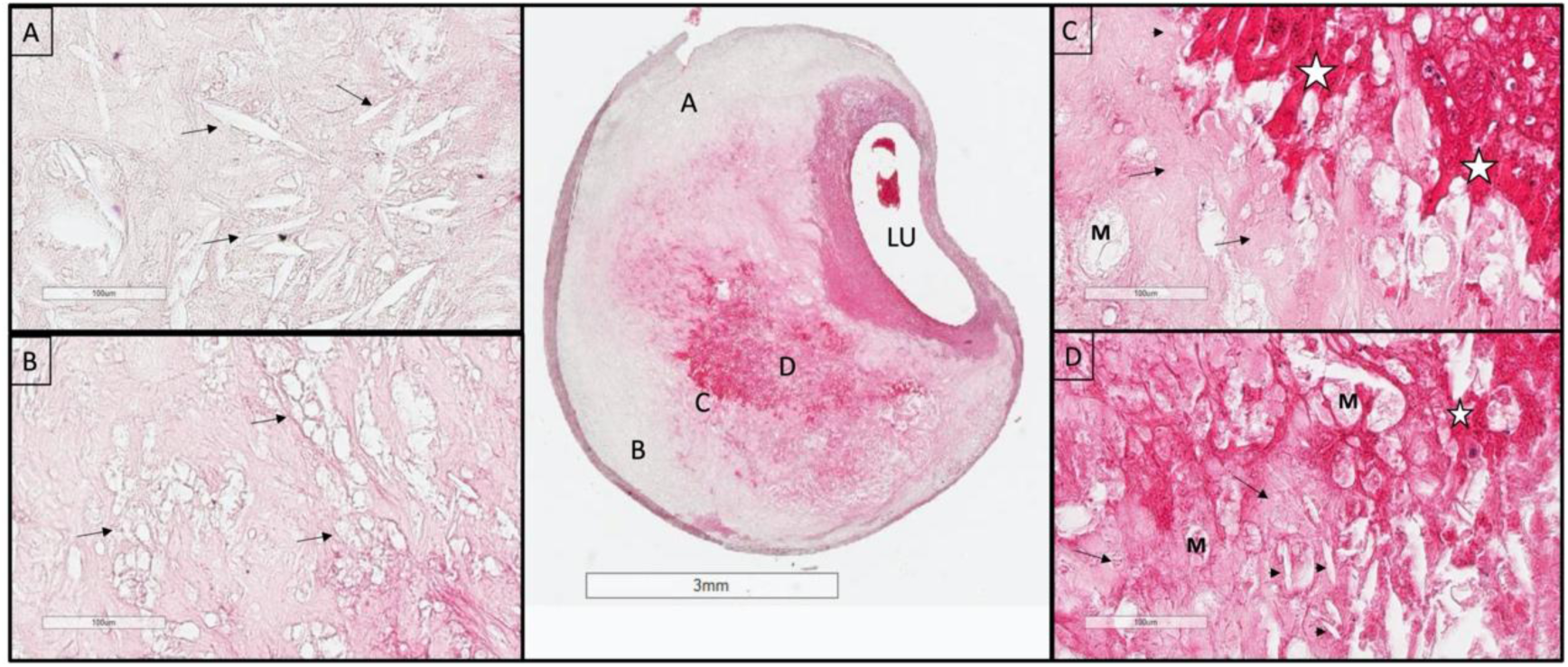
An example of the homogeneity of a Chinese plaque. Large deposits of minimally staining lipid and necrosis are predominant. A. Shows cholesterol clefts (arrows) in an acellular necrotic deposit. B. Shows clusters of dead macrophages (arrows) suspended in a simple matrix. C. Is taken at the junction of matrix (arrows) and intraplaque hemorrhage (IPH) (star). D. Illustrates the mixture of IPH (star), matrix (arrows), macrophages (M), and cholesterol clefts (arrow heads). LU indicates the lumen. (H&E)

**Figure 3.**
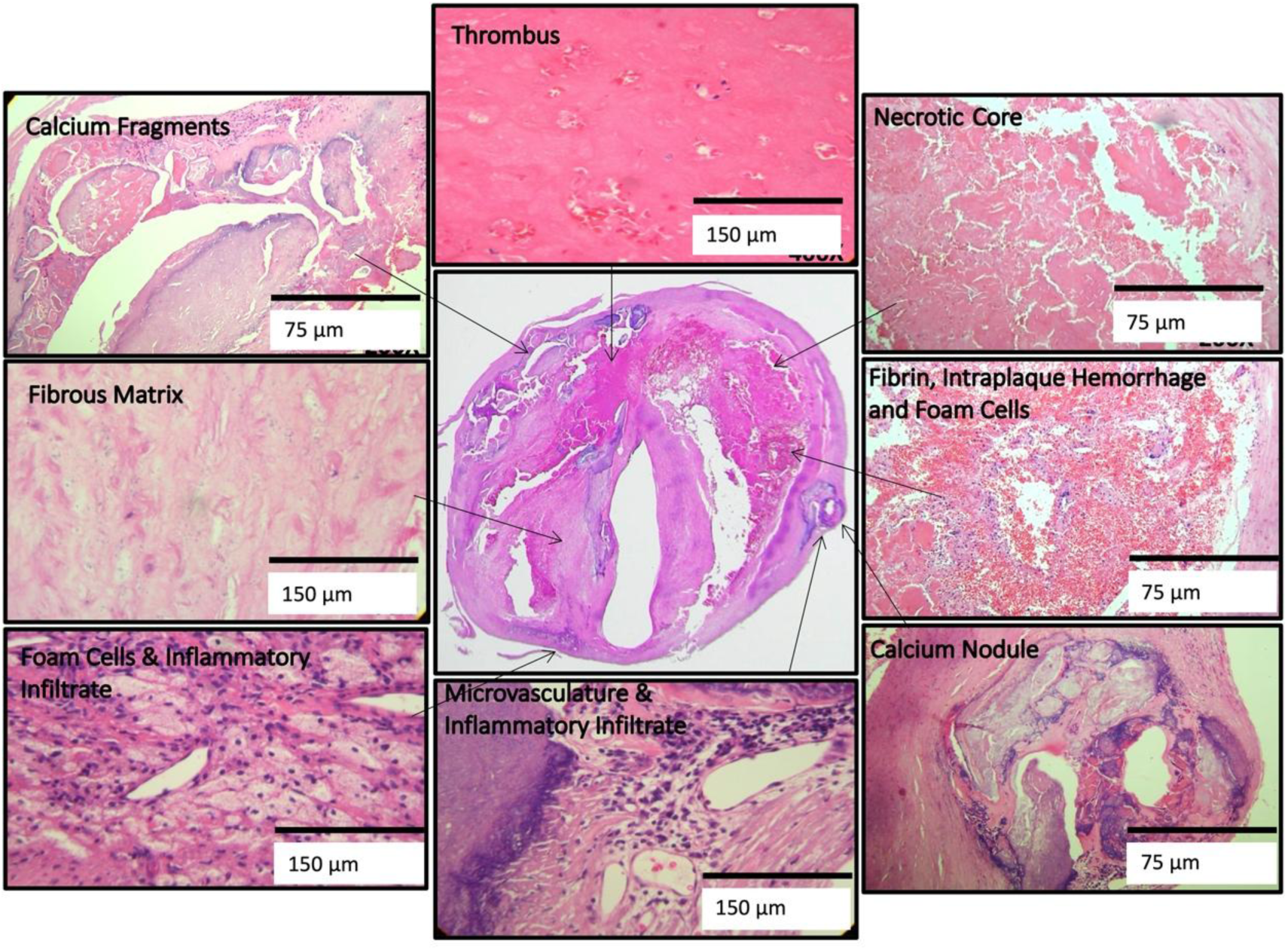
High power illustrations of common components found in the heterogeneous U.S. plaques (clockwise): thrombus; necrotic core; fibrin, intraplaque hemorrhage, and foam cells; calcium nodule; microvasculature and inflammatory infiltrate; foam cell aggregates and inflammation; fibrous matrix and calcium fragments. (H&E)

**Figure 4.**
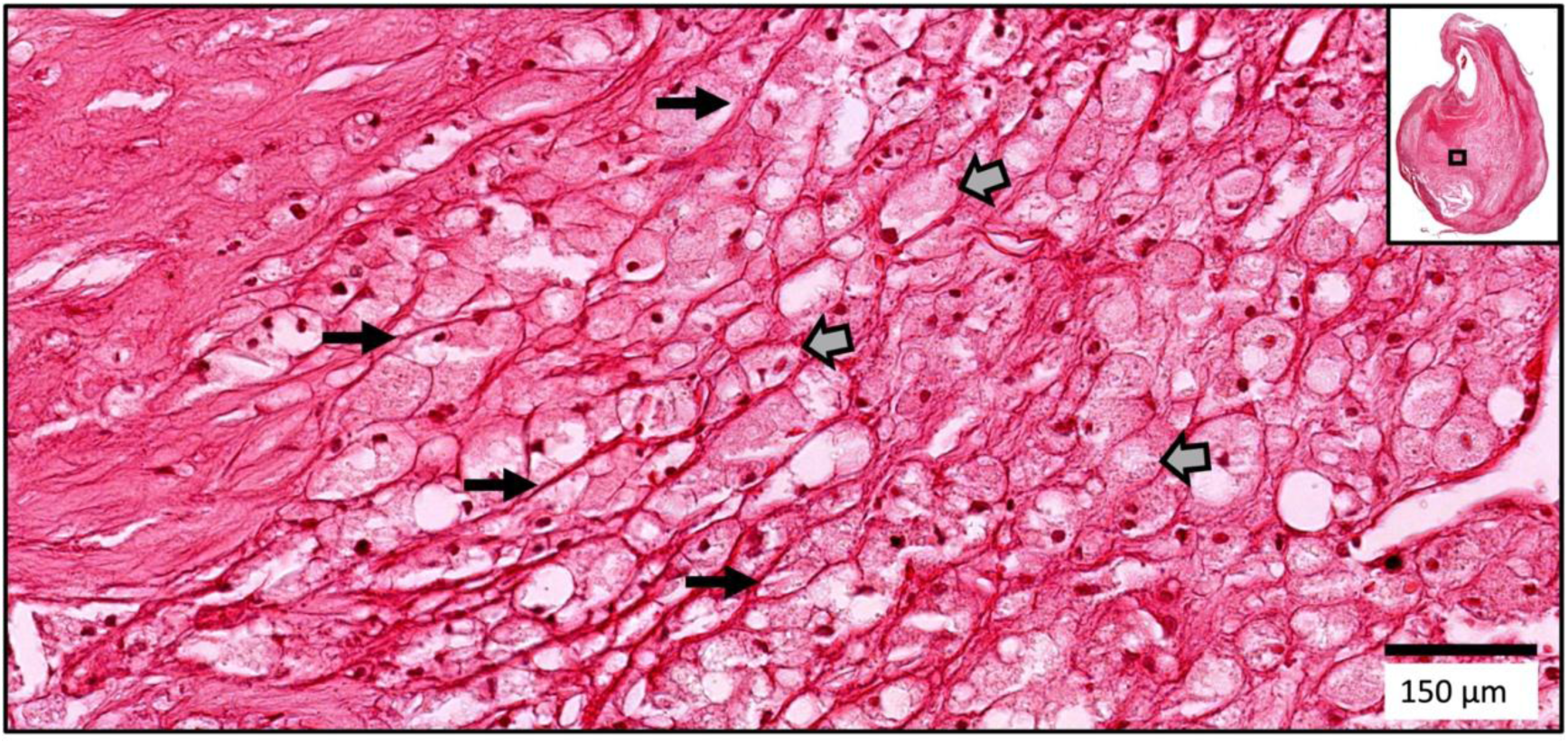
A fibro-lipid Chinese plaque. Large foam cells (block arrows) are sequestered along extracellular matrix fibers (long arrows) in this fibro-lipid Chinese plaque. (H&E)

**Figure 5.**
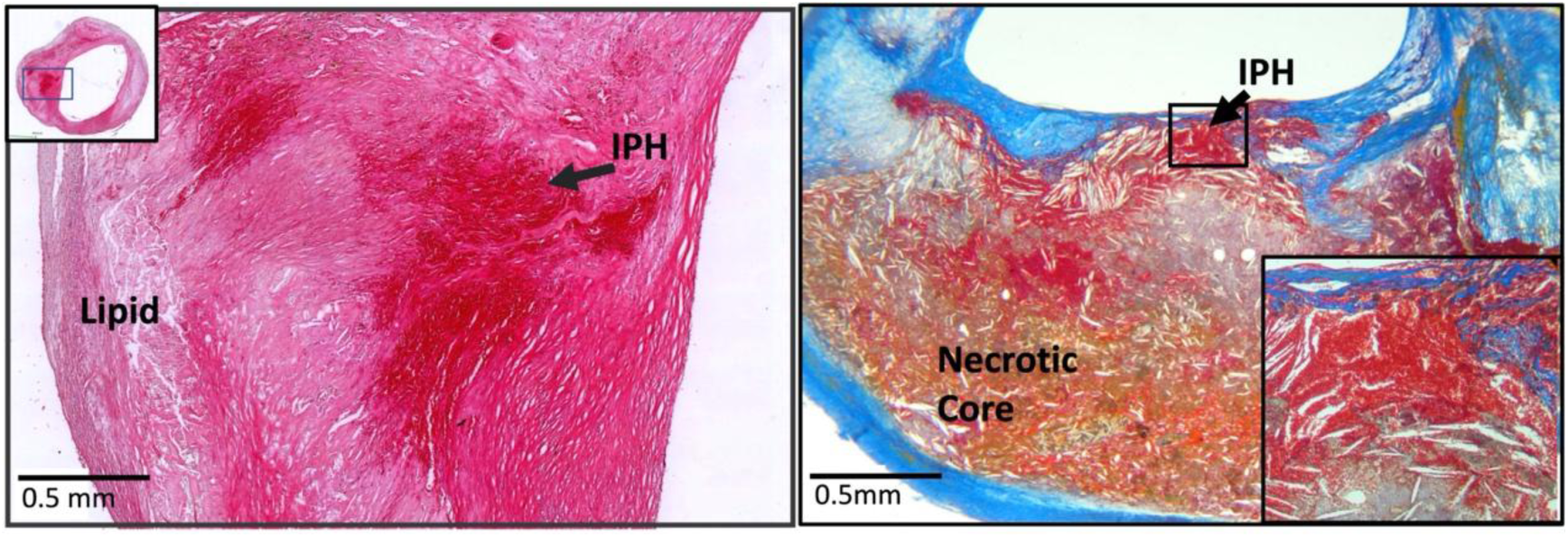
Recent intraplaque hemorrhage (IPH) into the fibrous matrix of a Chinese plaque (left) (H&E). Recent intraplaque hemorrhage into a necrotic core of a US plaque (right) (Mallory’s Trichrome).

Table 3 lists the difference in plaque composition between Chinese and U.S. groups. Prior to adjustment for possible confounders, the Chinese plaques were comprised of more lipid pools (11.6% vs. 2.9%, p<0.001) but fewer necrotic cores than the U.S. plaques (8.5% vs. 15.0%, p<0.001). The U.S. plaques contained more IPH overall than the Chinese plaques (17.7% vs. 9.3%, p=0.011). The Chinese plaques contained more recent IPH (5.2% vs. 1.5%, p=0.002) and less late IPH (2.0% vs. 10.2%, p<0.001) than the U.S. plaques. Volumes of calcification (median: 6.2% vs. 7.6%, p=0.63) and loose matrix (median: 1.2% vs. 1.4%, p=0.96) were similar in the two cohorts.

**Table 3.** Comparison of plaque composition and surface status.

| Composition Volume% | Cohort* |  |  | Adjusted Comparisons |  |  |
| --- | --- | --- | --- | --- | --- | --- |
| | Chinese group<br>(N=75) | U.S. group<br>(N=111) | P-value† | Adj.<br>$\beta$‡ | (95% CI) | P-value |
| Lipid pool, % | 11.6 (4.8 - 27.2) | 2.9 (0.7 - 9.1) | <0.001 | 10.0% | (4.9, 15.9) | <0.001 |
| Necrotic core, % | 8.5 (2.7 - 19.3) | 15.0 (4.1 - 26.1) | 0.031 | 0.3% | (-4.5, 4.9) | 0.94 |
| IPH, % | 9.3 (4.9 - 21.5) | 17.7 (6.9 - 28.0) | 0.011 | 0.2% | (-4.5, 4.7) | 0.94 |
| Recent IPH, % | 5.2 (0.9 - 17.4) | 1.5 (0.0 - 8.1) | 0.002 | 8.4% | (4.5, 12.7) | <0.001 |
| Late IPH, % | 2.0 (0.5 - 5.3) | 10.2 (2.4 - 21.9) | <0.001 | -8.2% | (-11.3, -5.4) | <0.001 |
| Calcification, % | 6.2 (2.5 - 15.9) | 7.6 (2.0 - 19.7) | 0.63 | 3.0% | (-3.1, 9.2) | 0.34 |
| Surface status |  |  |  | Adj.<br>OR‡ | (95% CI) | P-value |
| Fibrous cap rupture | 34 (45.3) | 74 (66.7) | 0.004 | 0.43 | (0.18, 1.04) | 0.061 |
| Mural thrombus | 10 (13.3) | 15 (13.5) | >0.99 | 0.45 | (0.11, 1.75) | 0.25 |
| Juxtaluminal calcification | 30 (40.0) | 65 (58.6) | 0.017 | 0.97 | (0.42, 2.27) | 0.95 |

OR = odds ratio for surface status; CI = confidence interval.
\*Values no. (%) or median (inter-quartile range).
†Fisher's exact test or the Wilcoxon rank-sum test comparing cohorts without covariate adjustment.
‡The regression coefficient ( $\beta$ ) or OR corresponds to the mean difference in composition or difference in risk of surface condition between plaques from the Chinese and U.S. cohorts after adjusting for symptom status, stenosis, sex, age, and the following risk factors and medications: hypertension, hyperlipidemia, diabetes, smoking status, coronary artery disease, statins, and use of antiplatelets/anticoagulants.

After adjustment for symptom status, stenosis, sex, age, and the risk factors and medications, Chinese plaques still had significantly greater volumes of lipid pools (adjusted β: 10.0%, [95% CI: 4.9, 15.9%], p<0.001) and recent hemorrhage (adjusted β: 8.4%, [95% CI: 4.5-12.7%], p<0.001) and smaller volumes of late hemorrhage than U.S. plaques (adjusted β: −8.2%, [95% CI: −11.3, −5.4%], p<0.001) (Table 3). However, adjusted differences in necrotic core (adjusted β: 0.3%, [95% CI: −4.5, 4.9%], p=0.94) and total hemorrhage (adjusted β: 0.2%, [95% CI: −4.5, 4.7%], p=0.94) were no longer statistically significant.

### Lumen Surface Status

Surface disruptions due to fibrous cap rupture were less common in Chinese specimens than in those in the U.S. cohort (45.3% vs. 66.7%, p=0.004) as were disruptions with juxtaluminal calcification (40.0% vs 58.6%, p=0.017). Frequencies of disruptions associated with mural thrombus were similar in the two cohorts (13.3% vs. 13.5%, p>0.99).

After adjustment for symptom status, stenosis, sex, age, the risk factors and medications, there was still a trend towards a lower prevalence of fibrous cap rupture in Chinese plaques than U.S. plaques (adjusted OR: 0.43, [95% CI: 0.18, 1.04], p=0.061). The prevalence of disruption associated with mural thrombus (p=0.25) or juxtaluminal calcification (p=0.95) was not significantly different between Chinese and U.S. plaques after adjustment.

### Associations of Plaque Features with Medications and Symptoms

After adjusting for cohort (Chinese vs. U.S.), symptom status, stenosis and other characteristics, there were no significant associations between statin use and either plaque composition or surface status (Table 4). IPH volume was significantly larger in plaques from patients with exposure to antiplatelet or anticoagulant agents than in those without this exposure (adjusted β: 4.1%, [95% CI: 0.0, 8.3%]. This difference in total IPH appeared mainly to be due to differences in recent IPH (adjusted β: 3.0%, p=0.059) rather than late IPH (adjusted β: 1.1%, p=0.47). There was also a trend towards an association between antiplatelet or anticoagulant use and juxta-luminal calcification (adjusted OR: 1.92, p=0.077).

**Table 4.**
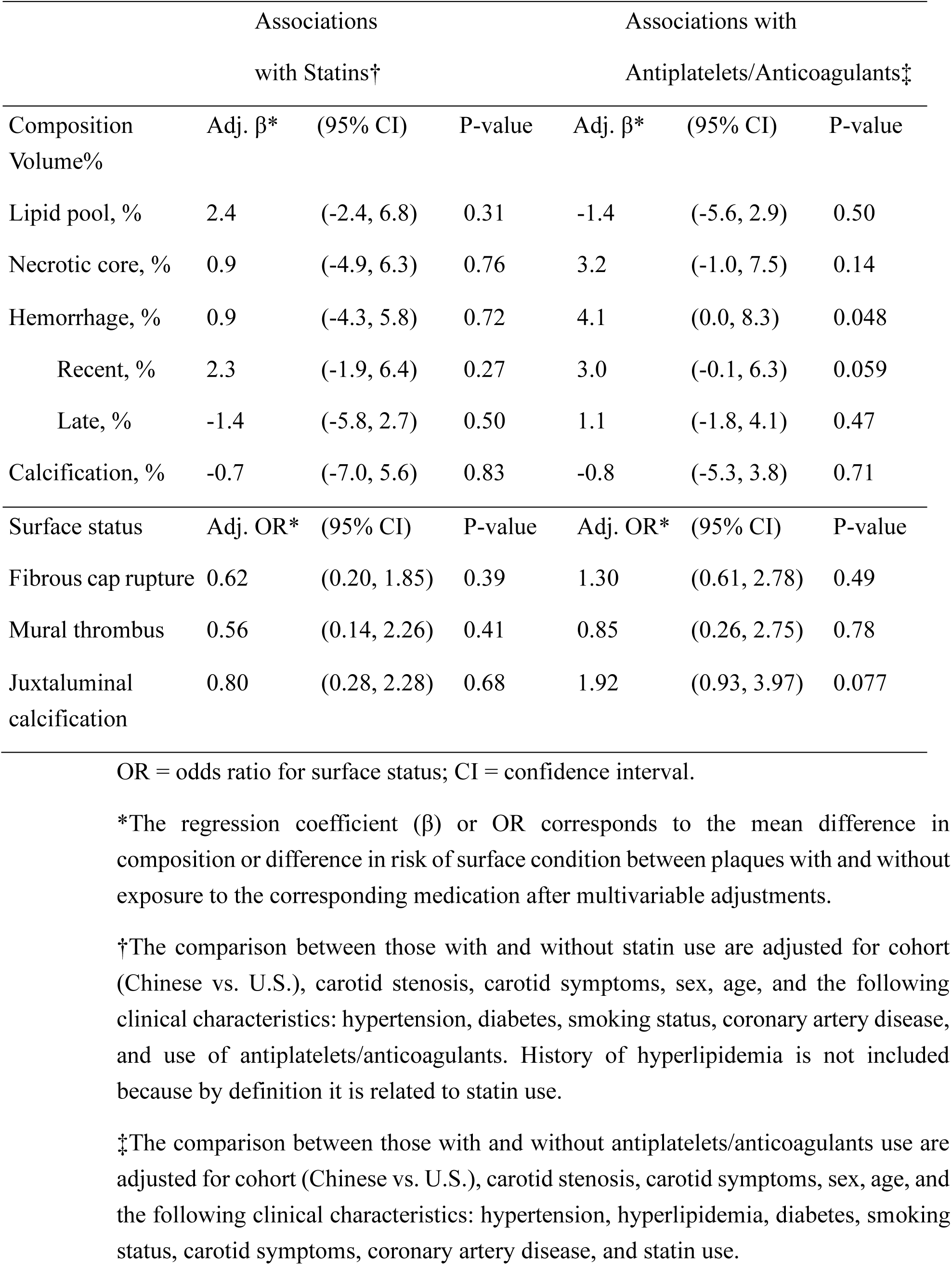
Associations between medications, plaque composition and surface status.

After adjusting for cohort, stenosis and other characteristics, there were no statistically significant associations between any plaque component or surface condition and carotid symptoms (Table 5). However, there were trends towards larger necrotic core (adjusted OR: 1.38 per 1-SD increase, p=0.096) and IPH (adjusted OR: 1.49 per 1-SD increase, p=0.052) being associated with symptoms. Recent IPH (adjusted OR: 1.33, p=0.13) and late IPH (adjusted OR: 1.31, p=0.18) had similar contributions. No associations with symptoms were significantly different between Chinese and U.S. plaques (p>0.14, Table 5).

**Table 5.** Associations of plaque composition and surface status with carotid symptoms.

|  |  | Associations<br>with Symptoms |  | Difference<br>between<br>Chinese and<br>U.S. |
| --- | --- | --- | --- | --- |
| % Composition Volume | Adj. OR* | (95% CI) | P-value | P-value† |
| Lipid pool, % | 1.16 | (0.80, 1.69) | 0.42 | 0.17 |
| Necrotic core, % | 1.38 | (0.94, 2.03) | 0.096 | 0.94 |
| Hemorrhage, % | 1.49 | (1.00, 2.22) | 0.052 | 0.77 |
| Recent, % | 1.33 | (0.92, 1.93) | 0.13 | 0.49 |
| Late, % | 1.31 | (0.89, 1.94) | 0.18 | 0.46 |
| Calcification, % | 1.10 | (0.79, 1.54) | 0.57 | 0.39 |
| Surface status |  |  |  |  |
| Fibrous cap rupture | 1.08 | (0.54, 2.16) | 0.83 | 0.63 |
| Mural thrombus | 1.55 | (0.60, 4.03) | 0.37 | 0.14 |
| Juxtaluminal<br>calcification | 1.32 | (0.66, 2.62) | 0.43 | 0.66 |

OR = odds ratio for association with symptoms; CI = confidence interval.
\*The OR for each composition variable is scaled to correspond to change per 1-SD increase after square root transformation to reduce skewness; the ORs are adjusted for cohort (Chinese vs. U.S.), carotid stenosis, sex, age, and the following clinical characteristics: hypertension, hyperlipidemia, diabetes, smoking status, coronary artery disease, statin use, and use of antiplatelets/anticoagulants.
†Wald test of an interaction between cohort (Chinese vs. U.S.) and the corresponding composition or surface status variable.

## Discussion

In the current study, the histological features of carotid plaque samples from China and the U.S. were comprehensively compared, with analysis of the entire length of CEA specimens removed without surgical disruption of the plaque or luminal surface. To our knowledge, this is the first such report incorporating detailed analysis of histological cross-sections obtained every 0.5 to 1.0 mm, thus avoiding bias from review of a smaller number of selected locations within the atheroma.

### Differences in Plaque Composition

Results demonstrated that there were significant differences in carotid plaque composition between the two groups. Despite more severe stenosis and larger plaque burden, Chinese plaques had fewer features of advanced atherosclerosis and were more homogenous, with significantly more recent IPH and larger accumulations of lipid. Late IPH and a trend toward more fibrous cap rupture, commonly considered as complex manifestations of a later stage of atherosclerosis, were found more commonly in U.S. plaques. These findings suggest that the fundamental pathophysiology of atherosclerosis in Chinese and U.S. cohorts may differ, and provides a histological reference for future comparative research, for example, in studies utilizing vessel wall MRI.^8^ Such studies, incorporating serial imaging of the carotid atheroma, will be critical to assess differences in response to treatment between groups, and perhaps aid in therapeutic decision making.

In a pilot study using MRI, plaque composition was compared between 20 symptomatic Chinese and 20 U.S. Caucasian patients with >50% stenosis.^7^ The U.S. cohort had a significantly higher prevalence of advanced lesions (AHA type VII) including luminal calcifications with juxtaluminal IPH and mural thrombus. Similar findings were noted in another MRI study of recently symptomatic Chinese and U.S. patients regardless of degree of stenosis: features of high-risk plaques (IPH or surface disruption) were significantly more frequent in extracranial carotids in the Caucasian cohort (42% vs 16%).^8^ These findings, along with dissimilarities in morphology and distribution noted in prior imaging studies, indicate significant differences in cerebrovascular pathophysiology between Chinese and North American populations, suggesting that a different approach to treatment and prevention may be needed.

### Difference in IPH Stage and Location

The detailed histological analysis of the specimens in this study identified distinct patterns of IPH in our two cohorts. Over 90% of both U.S. and Chinese plaques in our study had IPH. This high prevalence is consistent with the work of Derksen et al, who found an 81% prevalence of IPH in 794 carotid specimens.^18^ However, our histology results revealed that the stage and location of hemorrhage differed in our two cohorts. Recent IPH was present more frequently and in greater volume in Chinese plaques, while older IPH was more common and voluminous in U.S. plaques, suggesting that recurrent IPH over time into advanced plaques is more common in U.S. than in Chinese patients. This finding provides further evidence of the greater complexity of plaques in U.S. group. In U.S. plaques, almost all IPH occurred in the necrotic core, while in Chinese plaques, recent IPH was often found in fibrous matrix or adjacent to xanthomas or inflammatory infiltrate. The location of IPH (within a necrotic core or within fibrous matrix) might be a critical factor in how it impacts vulnerability to plaque rupture and embolization. Thus, further studies are needed to assess the association of recent IPH and its location to risk of embolic disease.

### Lipid Accumulation Patterns in Chinese Plaques

Results from this study revealed that Chinese plaques were more homogeneous with larger lipid accumulations. Further, a histologic feature identified as xanthomas was found much more commonly in Chinese plaques. The term xanthoma was utilized due to the similar histologic features as in xanthomas found in skin and tendon lesions. Morphologically, these distinct collections of foam cells embedded in a delicate and vascularized stroma resemble cholesterol--rich tendon xanthomas found in patients with familial hypercholesterolemia.^24^ This is the first study proposing this term with specific features in carotid plaques, which may be a consequence of most previous investigations being based on Western cohorts. Further studies are needed to ascertain the clinical significance and implications of carotid xanthoma. The differences in composition and complexity found in this study highlights the need for histological studies in populations with greater diversity in ethnic background and geography.

### Plaque Composition, Clinical Risk Factors and Medications

Our results showed that several clinical risk factors, as well as statin and anti-thrombotic drug use were significantly different between the two groups, which could be a reason causing the variations in carotid plaques. Hyperlipidemia was more common, and the percentage of statin usage was higher in the U.S. cohort. However, after adjusting for cohort (Chinese vs. U.S.), symptom status, stenosis and other characteristics, there were no significant associations between statin use and plaque composition. Antiplatelet use was associated with a greater volume of IPH, consistent with prior reports of increased IPH in association with the use of antithrombotic treatment.^25^ We found a higher prevalence of recent intraplaque hemorrhage in Chinese plaques, yet paradoxically, antiplatelet or anticoagulant use on admission was much less common in the Chinese cohort. Further MRI studies on clinical and risk factors associated with the development of new or recurrent carotid IPH are ongoing.

### Limitations

Since the plaque specimen was obtained from patients undergoing CEA, our cohort consisted of patients with significant carotid stenosis, and thus does not provide data on the characteristics of plaques in patients with mild to moderate stenosis. Also, we focused on histologic features that may be localized and quantified by clinical imaging techniques, realizing that more detailed analysis of cell types and local biochemical milieu also play a critical role in establishing plaque vulnerability. Another issue was that the collection time of the plaques in two groups of this study was approximately one to two decades apart, with the Chinese cohort more contemporary. Some of the differences noted in this study may be related to changes in disease pattern and treatment over time. However, our recent comparison of morphological and compositional characteristics in carotid plaques of two Chinese cohorts at different eras (2002-2005 and 2012-2015) using MRI showed there was no significant difference in IPH during different eras,^26^ which suggest that the differences in the pattern and stage of IPH between cohorts in this study are not entirely due to difference in time period. The former work also indicated that plaque lipid content was lower amongst the more recent cohort. In this paper, we found there were more lipid pools in Chinese compared to U.S. plaques. Further comparative studies by serial imaging are needed to confirm the results of this study, and to determine the natural history and clinical outcome associated with various plaque morphologies.

## Conclusions

In conclusion, Chinese plaques had fewer features of advanced atherosclerosis and were more homogenous with significantly more recent IPH and larger accumulations of lipid than US plaques. These findings demonstrate that atherogenesis may produce fundamentally different lesions in different populations. Understanding the racial differences may allow better risk assessment and individualization of treatment in different populations.

## Clinical perspective

### Clinical Competencies

1. Carotid plaque composition was significantly different in these Chinese and U.S. cohorts.
2. Chinese plaques were less complex with larger lipid accumulations, more recent intraplaque hemorrhages, and tended to have fewer fibrous cap ruptures. U.S. plaques had more advanced features such as late intraplaque hemorrhage.

### Translational outlooks

1. These histologic findings together with prior MRI studies suggest a significant difference in the underlying pathophysiology of cerebrovascular disease in Chinese and U.S. populations that may require a different approach to management.
2. Investigations of the racial difference in carotid advanced atherosclerosis may provide insight into variation in response to therapy between ethnic groups and help guide strategies for treatment.
3. Further studies on intra- and extracranial atheroma with mild to moderate stenosis, along the follow-up imaging will provide insight into whether there are any differences in earlier stages of atherosclerosis development and progression between Chinese and U.S. cohorts.

## Acknowledgments

We would like to thank the individuals who volunteered to participate in this study.

## Conflict of Interest

All the authors listed have contributed to the manuscript and approved its final submitted form, and that the authors have read and agree to Editorial Policies. The authors report no conflicts of interest in this work.

## Abbreviations and Acronyms

AS: Atherosclerosis
MRI: magnetic resonance imaging;
CEA: carotid endarterectomy;
LRNC: lipid rich necrotic core;
IPH: intraplaque hemorrhage;
CI: confidence interval;
OR: odds ratio;
H&E: hematoxylin and eosin

---

**Supplement Figure 1.**
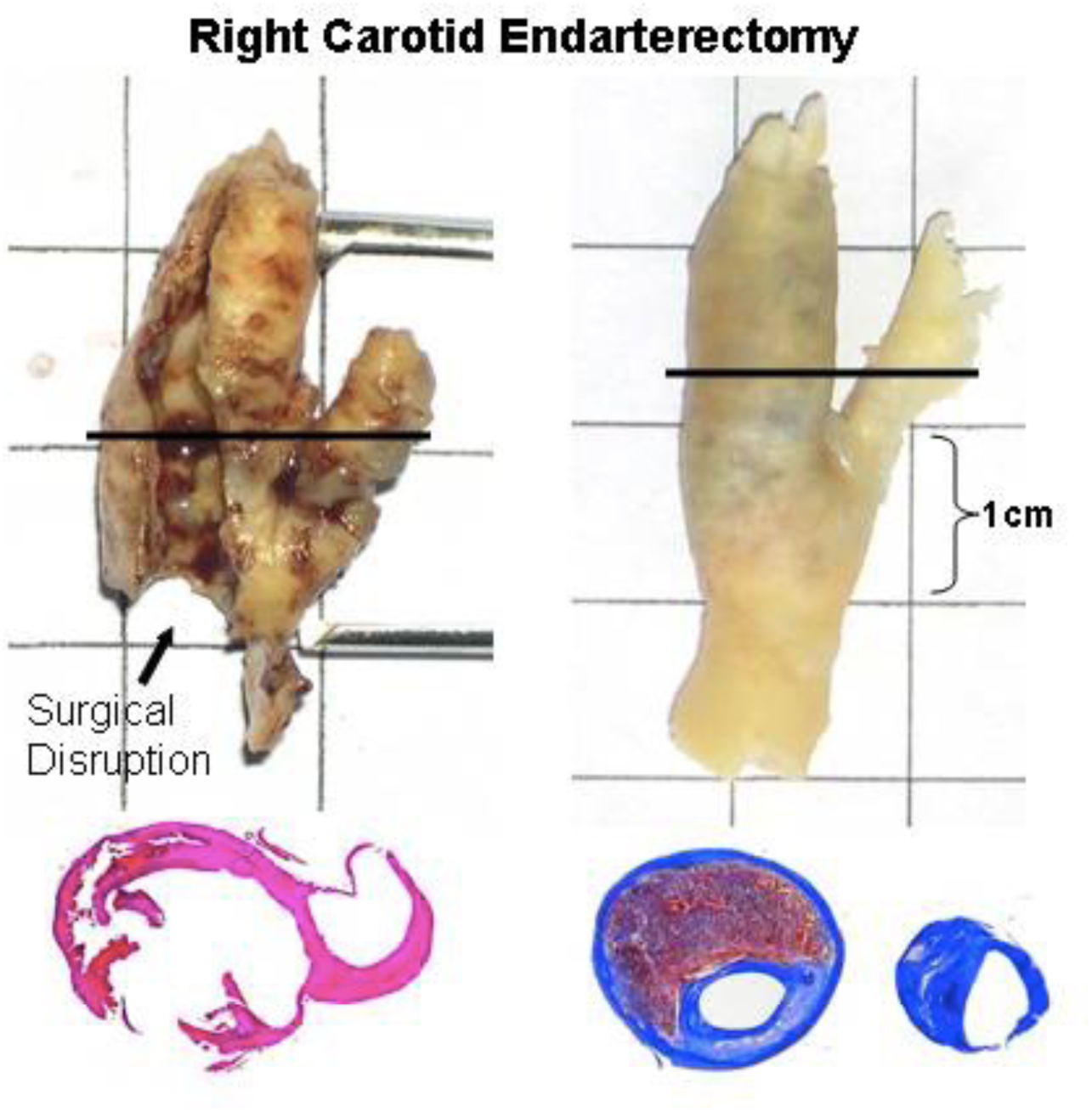
Left. Carotid plaque from a standard endarterectomy with disruption of the specimen. Right. Specimen from the modified surgical technique without disruption of the inner surface. Corresponding histologic sections are below.

**Supplement Figure 2.**
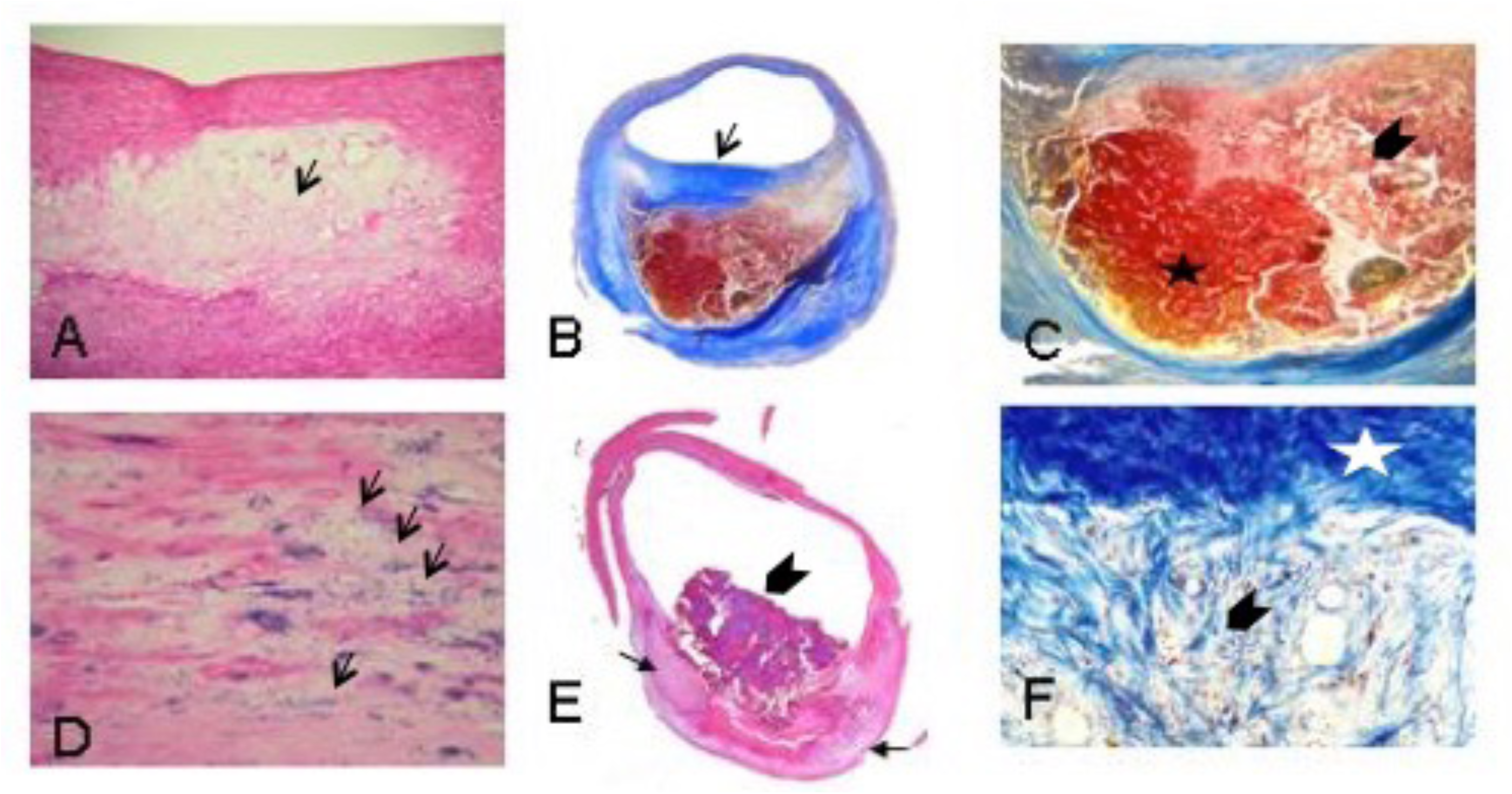
Carotid plaque components: A) Lipid pool (arrow)(H&E); B) Necrotic core with thick fibrous cap (arrow) (Mallory’s trichrome); C) Recent intraplaque hemorrhage into a necrotic core (star)(H&E). Late intraplaque hemorrhage in the necrotic core (chevron) (Mallory’s trichrome); D) Speckled calcification (arrows)(H&E); E) Protruding calcium nodule (chevron) with two calcified plates (arrows)(H&E); F) Loose matrix (chevron) with adjacent dark blue dense matrix (star) (Mallory’s trichrome).

**Supplement Table 1.** Correlation of plaque composition with stenosis and percent wall volume (N=186)

|  | Correlation with Stenosis |  |  | Correlation with Percent Wall Volume |  |
| --- | --- | --- | --- | --- | --- |
| Component | Spearman's r | p |  | Spearman's r | p |
| Lipid pool | 0.30 | <0.001 |  | 0.23 | 0.002 |
| Necrotic core | 0.30 | <0.001 |  | 0.34 | <0.001 |
| Hemorrhage, % | 0.29 | <0.001 |  | 0.39 | <0.001 |
| Recent hemorrhage, % | 0.30 | <0.001 |  | 0.35 | <0.001 |
| Late hemorrhage, % | 0.00 | 0.98 |  | 0.15 | 0.048 |
| Calcification, % | -0.25 | 0.001 |  | 0.06 | 0.44 |
| Loose matrix, % | 0.25 | 0.001 |  | 0.15 | 0.044 |

## Notes

**Disclosure of Conflict of Interest:** All the authors listed have contributed to the manuscript and approved its final submitted form, and that the authors have read and agree to Editorial Policies. The authors report no conflicts of interest in this work.

### Competing Interest Statement

The authors have declared no competing interest.

